# Characterization of structural and energetic differences between conformations of the SARS-CoV-2 spike protein

**DOI:** 10.1101/2020.11.01.363499

**Authors:** Rodrigo A. Moreira, Horacio V. Guzman, Subramanian Boopathi, Joseph L. Baker, Adolfo B. Poma

## Abstract

The novel coronavirus disease 2019 (COVID-19) pandemic has disrupted modern societies and their economies. The resurgence in COVID-19 cases as part of the second wave is observed across Europe and the Americas. The scientific response has enabled a complete structural characterization of the Severe Acute Respiratory Syndrome – novel Coronavirus 2 (SARS-CoV-2). Among the most relevant proteins required by the novel coronavirus to facilitate the cell entry mechanism is the spike protein trimer. This protein possesses a receptor-binding domain (RBD) that binds the cellular angiotensin-converting enzyme 2 (ACE2) and then triggers the fusion of viral and host cell membranes. In this regard, a comprehensive characterization of the structural stability of the spike protein is a crucial step to find new therapeutics to interrupt the process of recognition. On the other hand, it has been suggested the participation of more than one RBD as a possible mechanism to enhance cell entry. Here we discuss the protein structural stability based on the computational determination of the dynamic contact map and the energetic difference of the spike protein conformations via the mapping of the hydration free energy by the Poisson-Boltzmann method. We expect our result to foster the discussion of the number of RBD involved during recognition and the repurposing of new drugs to disable the recognition by discovering new hotspots for drug targets apart from the flexible loop in the RBD that binds the ACE2.

## INTRODUCTION

Each year numerous diseases are caused in humans by life-threatening viruses (as classified by the World Health Organization (WHO), e.g., the seasonal flu (influenza A and B), yellow fever, ebola, rabies, coronaviruses (CoVs), among others). Although CoVs were only associated with a common cold, for the first time in 2002, a new kind of CoV led to the medical condition known as the severe acute respiratory syndrome (SARS-CoV). This new disease was discovered in southern China and infected more than 8000 people in 27 countries, causing 774 deaths worldwide^1^. Later, in 2011 a more deadly coronavirus emerged in the middle east and was named the Middle East Respiratory Diseases (MERS) as it mostly affected people in Saudi Arabia (infected 4985 cases and 858 deaths) with a high fatality rate of 34% that was almost four times larger than the previous outbreak^2^. However, neither of those previous outbreaks stressed the worldwide health system and economy^3^ more than the novel coronavirus. The WHO has labeled the novel coronavirus as SARS-CoV-2 due to its structural similarity with SARS-CoV, and the disease caused by it as COVID-19. On March 11, 2020, the WHO officially declared the outbreak of COVID-19 as a pandemic.. As of October 30, 2020, almost 45M confirmed cases with a death toll over 1M around the world have been reported. Thus, there is an urgent need to understand the molecular features of each of the proteins that are assembled into the virion. The characterization of the conformational changes related to proteins that compose the structure of the virus and its connection with the high spreading of the disease are crucial steps, which in turn will permit the development of more effective therapeutics for fighting against the disease, early prevention as well as controlling future outbreaks.

The fast spread of COVID-19 around the globe has triggered the immediate response of the scientific community that aims to understand the pathogenesis of the novel coronavirus and in particular the molecular mechanisms of SARS-CoV-2 during cell entry which can aid to devise viral deactivation strategies prior to cell recognition or block the viral replication mechanism, among others^4^. A key component in all coronavirus associated with cell entry is the spike protein (S) and thus its structural characterization is an essential step^5,6^. The typical spike protein is a homotrimer system responsible for the “corona” (Latin word for crown) appearance in all coronaviruses, and it plays a crucial role in the interaction between the virion and the host human cell membranes. The spike protein attaches itself to specific cellular receptors (i.e., human angiotensin-converting enzyme–ACE2)^7,8^ and undergoes several conformational changes that engage different protein domains (e.g. receptor binding domain-(RBD), N-terminal domain-NTD, S1, and S2 subunits) (see Figure 1). The first process that is believed to occur relates to the transition from down to up receptor-binding domain conformation. This transition prepares the virus for binding to the ACE2 receptor and the subsequent internalization of the virus through the formation of the endosome, later fusion of the viral and cell membranes, and the final release of the viral RNA into the cytoplasm^9^. It is clear that the spike protein is a crucial component in each aspect of the cell entry mechanism. Several studies^5,10–14^ have elucidated how the novel coronavirus takes advantage of the spike protein structure to outperform SARS-CoV. For instance, the characterization of the mechanical stability of the RBD of SARS-CoV-2 has shown it to be stiffer (greater by 50 pN) compared to SARS-CoV^15^. This result has important consequences during binding ACE2 (pre-fusion state)^16^, as it can withstand Brownian and cellular forces and yet maintain close contact while priming of the spike protein by transmembrane protease serine 2 (TMPRSS2) occurs as part of S1 dissociation from S2 that enables the post-fusion mechanism^16,17^. In addition, *in silico* studies found a space correlation between the polybasic furin cleavage site Q_677_TNSPRRAR↓SV_687_ and surface residues located in the RBD region that recognize the ACE2 in SARS-CoV-2. Such effect was mediated by a long-range electrostatic interaction 10 nm apart^18^. Furthermore, the mutant D614G (a single aa change, D = aspartic acid by G = glycine) of the SARS-CoV-2 spike protein sequence, which began spreading in Europe in February and became the dominant form globally at the end of March, displayed stronger transmissibility^19^. Also, this mutation correlated residues that are located about 7–10 nm from the SARS-CoV-2 RBD^18^. Certainly, it has also been suggested that those features in the spike protein could enhance its transmissibility and facilitate the post-fusion machinery in SARS-CoV-2^20^. The intrinsic flexibility of the full ectodomain dictated by three hinges was characterized by cryo-EM and large-scale MD simulation^21^. It shows the flexibility of the spike protein and the ability of the spike head to explore different orientations in space which allows it to scan the host cell surface in search of ACE receptors. A recent cryo-EM study has found a free fatty acid pocket in each RBD^22^. The binding of the linoleic acid (LA) stabilizes a locked conformation giving rise to reduced ACE2 interaction in vitro. The sugar coating of the surface in the spike protein by glycans are not only shielding to evade the immune system response as commonly believed^23^, but also they may play a structural role by modulating the conformational dynamics of the spike’s RBD that is responsible for cell recognition^24^.

**Figure 1:**
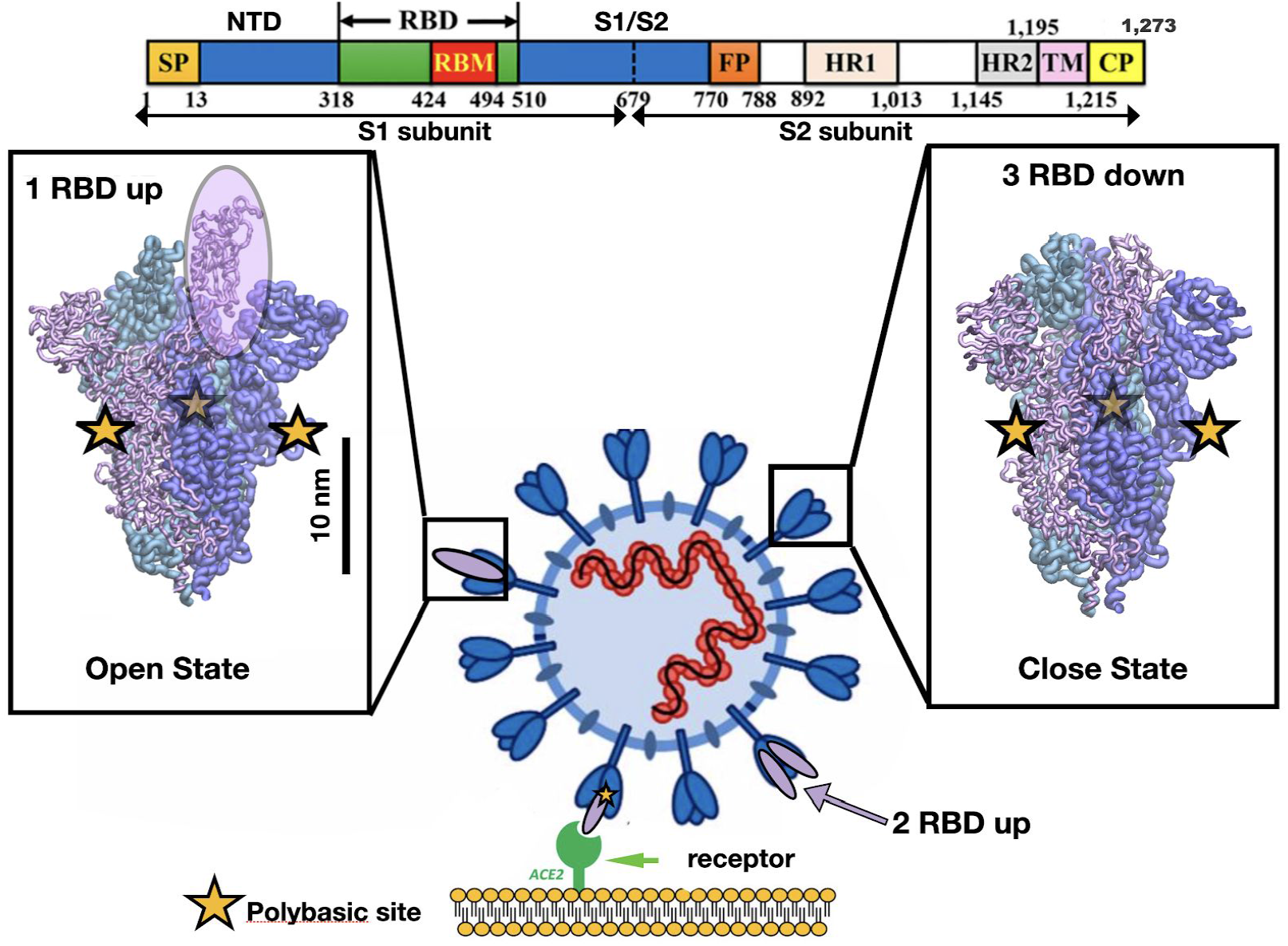
Representation of the different conformations of the RBD in the SARS-CoV-2 spike protein. Cell recognition is initiated by the RBD transition from down to up conformation and then the high affinity triggers binding between the RBD in the up conformation and the ACE2 receptor shown in purple and green respectively. The sequence of one chain of the spike protein is shown on top as well as the residue numbers for several protein domains. The bar shows the typical length scale for the whole system. The fusion of the viral and cell membrane takes place by surface proteases that cleave each chain at the polybasic sites (yellow stars) located at the interface of S1/S2 subunits.

In order to fight against COVID-19 several medical strategies are developed by targeting the SARS-CoV-2 spike protein such as the development of viral vector and protein-based vaccines^25^ and monoclonal antibodies treatment such as regeneron cocktail that blocks the virion by binding to RBD^26,27^. New efforts in novel therapeutics employing short peptides, proteins, and natural resources such as plant derivatives are yet new ways for disabling the virion at the level of the spike protein^28–31^. In this regard, a stabilized form of spike protein with all RBD in down or one (or two) RBD in the up conformation is desirable for vaccine and therapeutic development because this conformation displays most of the neutralizing epitopes that can be targeted by antibodies to prevent cell entry^6,32^. These stabilized structures contain two consecutive proline substitutions in the S2 subunit in a turn between the central helix (CH) and heptad repeat 1 (HR1) that is important during the transition to a single, elongated α-helix in the postfusion conformation. Prefusion-stabilized spikes in closed and open states have been used to determine high-resolution spike structures by cryo-EM^5,7^ that has been crucial for large-scale MD simulation restricted to microsecond time scale. However, even with these substitutions, the SARS-CoV-2 ectodomain is unstable without the ACE2 receptor and typically difficult to express on mammalian cells, hampering biochemical research and search for novel vaccines. A recent smFRET study characterizes the ensemble of conformations occurring in the spike which seems to be part of a dynamic equilibrium^33^, but it does not lead to a quantitative assessment of the relative stability between conformers. A multiscale modeling that employs structure-based coarse-graining has shown the existence of a dynamic asymmetry that triggers the change in conformation of the closed to open states and the characteristic free-energy landscape shows the closed conformation as the ground state^33,34^. Here we show an analysis of the relative stability taking into account dynamic contact map analysis. Our study shows the correlation between different conformations of the spike protein and its RBD, NTD, and S2 subunits and highlights the role of destabilization in order to get access to other conformations. This tool also allows determination of stable “hotspots” across the amino acid sequences and possible new targets for therapeutics. In addition, we obtain free energy differences between states of the spike protein without ACE2.

## MATERIALS AND METHODS

### Modeling of the full SARS-CoV-2 spike structures

The SARS-CoV-2 spike (S) protein conformations in the closed and open states were modeled with all RBD in down conformation and with one or two RBD in the up position. The different conformations studies correspond to three RBD in the down position (3down), one RBD in the up and two RBD in the down positions (1up2down) and two RBD in the up and one RBD in the down position (2up1down). They were reconstructed based on cryo-EM structures with PDB codes 6VXX, 6VSB and 6X2B respectively. The wild-type (WT) sequences that describe the spike protein come from QIQ50172.1 stored in the GenBank database for SARS-CoV-2. The trimeric cryo-EM structures comprise mainly the sequence A27-S1147. In order to stabilize the cryoEM pre-fusion state several mutations were implemented, e.g. P986K and P987V^5^. In structure files (PDB), however, there are missing residues that must be fulfilled to get the correct WT model, as well as, some residues that must be mutated to reconstruct the WT type. In particular, using as reference QIQ50172.1 starting at residue MET 1, residues ALA 570, THR 572, GLN 607, GLY 614, ARG 682, ARG 683, ARG 685, PHE 855, ASN 856, LYS 986, and VAL 987 were replaced from original PDB structures to reconstruct the WT sequence. The standard Needleman–Wunsch algorithm was used as implemented by Chimera visualization software to align the sequences and the missing loops were modeled by homology using MODELLER^35^ and a few steps of energy minimization algorithms. The disulfide bonds were the ones prescribed by the PDB files and 14 per single chain of the spike homotrimer.

### All-atom MD of the SARS-CoV-2 Spike and its conformations

Amber18^36^ was used to carry out all-atom simulations. The protein, water, and ions were all modeled using the FF14SB^37^ and TIP3P^37,38^ force fields. System energy was minimized using the CPU version of pmemd, while heating, equilibration, and production simulation stages used GPU pmemd. SARS-CoV-2 systems 3down, 1up2down, and 2up1down were solvated in octahedral water shells of 14 Å, 12 Å, and 16 Å, respectively. Disulfide bonds (DBs) were added between cysteine residues identified in the initial models as being close enough for a DB bond to form. The DBs were added using tLeap. The NaCl concentration for all simulations was 0.150 M NaCl. In order to use a 4 fs time step, the hydrogen mass repartitioning was applied to the protein^39^. SHAKE was applied to hydrogens, and an 8 Å real-space cutoff was used. For long-range electrostatics we utilized PME with periodic boundary conditions. Minimization included 2000 steps of steepest descent followed by 3000 steps of conjugate gradient method. The heating protocol used: (1) Heating from 0 to 100 K (50 ps) in NVT, and (2) heating from 100 to 300 K (100 ps) in NPT. During minimization and heating restraints of 10 kcal mol^-1^ Å^-2^ were applied to C_α_ atoms. Subsequently, equilibration at 300 K was carried out and C_α_ restraints were gradually reduced from 10 kcal mol^-1^ Å^-2^ to 0.1 kcal mol^-1^ Å^-2^ over 6 ns. For the production simulations, restraints were released and 200 ns and 320 ns unrestrained production simulations were carried out for 3down and 1up2down SARS-CoV-2 conformations respectively and 100 ns for 2 up 1 down case. Production simulations started from the final equilibrated coordinates, and for each system five replicas were simulated. Pressure of 1 atm was maintained using the Monte Carlo barostat, and a temperature of 300 K was maintained during production using the Langevin thermostat (collision frequency 1 ps^-1^), as implemented in Amber18^36^. In aggregate, 2.3 μs of all-atom MD simulation data was used for this work. Snapshots from the MD simulations can be found in a Zenodo repository^40^.

### Differential Contact Map (dCM) analysis

The contact map (CM) determination considers the VdW interaction between residues that are typically captured by a geometric based-approach denoted by extended overlap (OV) of the VdW spheres, which has been successfully used before to describe single proteins^41–44^. Furthermore, the chemical character of the residues can be also included as an additional part of the CM determination. The latter is denoted as the rCSU approach, which considers the chemical character of each atom, and respective possible bonds between two residues, into categories that count the number of stabilizing and destabilizing contacts per residue, defining a contact when both residues have a net stabilizing character. Together they form a robust CM methodology known as OV+rCSU contact map^45^ that has been validated in the dynamic of large protein complexes^46–50^. This approach is used to get structural information from a specific geometry. This methodology is reliable enough to describe relatively small globular protein molecules. However, when applying it at a larger complex system, we may include contacts between residues that are not relevant to the system, e.g., contacts that belong to solvent-exposed flexible loops are less structurally relevant to describe a protein than the ones between residues of α-helices and β-strands. To get only the most structural relevant contacts, we can use the information available from the molecule’s dynamics. Computing the contact maps for each frame from a subset of MD frames, we can count how many times a given contact was identified. Then, a contact has a *frequency*, freq, defined by the number of frames where it was found over the total number of frames analyzed. This procedure gives a low frequency for contacts between flexible parts of the protein, while highly stable structures exhibit high frequency.

Here, we use this methodology to sweep evenly distributed frames of the equilibrium MD trajectory of each system studied in this article to dynamically determine the high frequency contacts (freq> 0.9) between amino acids. In particular, from 10,000 frames of the closed conformation, we obtained a total number of 29,334 ± 82 contacts, 8,000 frames for the 1up2down conformation showing 29,320 ± 741 contacts, as well as 8,340 frames for 2up1down case with 29,055 ± 718 contacts, which is a robust standard deviation of less than 3% from the total number of contacts. The source of contact fluctuations are essentially flexible loops that account for approximately 1772 amino acids based on secondary structural analysis, while helices and strands are approximately represented by 712 and 819 residues, respectively. The whole spike protein has a total of 3363 residues. Our objective is to discern between relevant and not relevant contacts, and a moving coil that eventually creates a contact is obviously less stable than a secondary structure. These small deviations in the number of contacts, and the consequent contacts frequencies, allow us to differentiate between relevant and not relevant contacts for the system’s structural stability, which is the main advantage of this methodology compared to a static analysis based on only one frame mostly taken from X-ray/NMR crystallography. To be able to compute such a large amount of contact maps, we implemented our own contact map software that implements the OV+rCSU approach, as detailed in Ref.^45^, and available via Zenodo repository^40^.

### Poisson Boltzmann calculations for the spike protein energetic characterization

Rather than considering water molecules explicitly, an implicit solvent model averages their influence in a continuum dielectric description^51^. Then, a dissolved molecule is represented as a multi-dielectric infinite domain with, at least, two regions ---the solute and the solvent---interfaced by the solvent-excluded surface (SES). This situation in the case of the spike proteins is divided in two regions:

- Coronavirus spike protein structures (**Ω**_1_).
- External (**Ω**_2_), representing the solver.

Each region is characterized by its dielectric constant (**ε**_1_, **ε**_2_), and salt concentration (***C_1_, C_2_***), which is considered as zero in the solute. For simplicity, we will use parameters that are proper for a solvent in the external region (**Ω**_2_).

The partial charges in the solute are represented as static point charges at the locations of the atoms, whereas the salt ions in the solvent are considered as mobile point charges that arrange according to Boltzmann statistics. Applying continuum electrostatic theory on this arrangement gives rise to a system of coupled partial differential equations, where the potential inside the solute is governed by Poisson, and in the solvent by Poisson-Boltzmann:

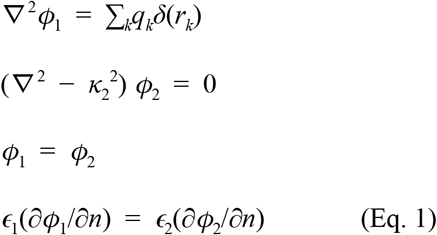

Here, **Φ** is the electrostatic potential, **κ** is the inverse of the Debye length (which depends on the salt concentration ***C***), **ε** is the dielectric constant, and ***q*** the partial charges of the biomolecule. **Γ**_1_ is the SES on the outside of the Spike protein, interfacing **Ω**_1_ with **Ω**_2_. The unit vector ***n*** is normal to the SES, and points away from the region enclosed by the surface.

The most common quantity of interest is the solvation energy, which is the work required to bring the solute from vacuum into the solvent. Considering that the charges inside the solute are Dirac delta functions, this energy becomes

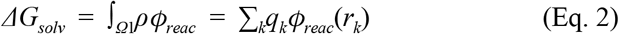

where *ϕ_reac_* = *ϕ* – *ϕ_coul_* is the reaction potential and **ρ** is the charge distribution, and ***r***_*k*_ the location of charge *k*. There is another source of energy in the point-charge distribution of the partial charges in the solute. Then, the total electrostatic contribution to free energy is the sum of the solvation energy in Eq. 2 and the Coulomb-type energy

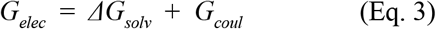

where,

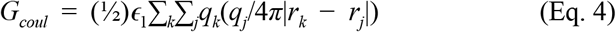

In this work, we use the Poisson-Boltzmann solver PyGBe^52^ to compute the electrostatic potential and energy in equations Eq. 1 and Eq. 3. PyGBe formulates the system of partial differential equations in a boundary integral form, requiring to mesh the SES only, and solves the resulting system with a boundary element method (BEM)^53^. From a viewpoint of MD simulation trajectories, we based the PB initial structures to be meshed on the final snapshots of the MD trajectories for the three studied spike proteins. In addition to the structural positions the charges and vdW radii are also required by the PB solver to calculate solvation and Coulomb energies, mostly given in a pqr format. Here, we remark that PB-energies provide a static picture of the solvation and Coulomb energies at a certain point of the MD simulations trajectory. In this case we employed the final MD conformation snapshots of the different simulation replicas^40^ corresponding to the three systems under consideration.

## RESULTS

### Relative stability analysis of the SARS-CoV-2 spike protein conformation

The characterization of the full space of conformations of the spike protein and the intrinsic stability by MD dynamics simulation is still limited by the length and time scales needed to sample the conformational space of the spike protein. In a recent study carried out by smFRET^33^ the closed and open conformations were sampled by tagging certain amino acids in the NTD and RBD regions and monitoring their positions in space. In particular, this methodology allows the reconstruction of a restricted free energy landscape that shows a high stability for the closed state over the open, as it is believed in the absence of the ACE2 receptor. However, it does not highlight the amino acid regions or the hotspots of stability which can be used for therapeutics by developing small drugs that bind those regions and disrupt the stability associated with them. In MD simulation the transition from closed to open states is only possible via large forces acting on the RBD in a nonequilibrium fashion. This protocol captures the transitions but it is only thermodynamically consistent in the limit of a large number of pathways through the Jarzynski equality^51^. In this section we define the local stability of the homotrimer spike protein defined by the difference of the number of native contacts associated with single amino acids between two conformations. The change of the number of contacts under a conformational change shows a measure of the loss or gain of the local stability. We test our dCM methodology on three different conformations of the spike protein. As a result we report the number of destabilizing residues and contact set for each conformation in Table 1. Based on our analysis the 1up2down conformation shows less number of destabilizing residues than the 2 up 1 down conformation. However, the stabilizing aa residues per protomer in up conformation seems to be comparable. Here we prove that among the two studied conformations of the open state the most stable is the 1up2down system. In Table 1 we report the number of the destabilizing and stabilizing residues and their corresponding high-frequency (freq > 0.9) contacts. At first, we observe the large number of destabilizing residues 92 and 502 which account for 3% and 15% of the total system for 3down to 1up2down and 3down to 2up1down respectively. This is an indication that a transition from closed state to 1up2down implies the destabilization of a small number of native contacts. However, such a type of transition is more serious as it destabilizes several parts of the protein once it exposes 2 RBD in up conformation. Note that a small gain in stability is reported in very key places of the spike protein such as flexible loops that are engaged in cell recognition at the RBD (chain B) and stabilized by formation of new intrachain contacts. In the absence of ACE2 our study renders the open state less stable than the closed state which is in agreement with smFRET study^33^ that identifies the closed state as the most populated state in the lack of ACE2. Moreover, our study shows an additional structural stability associated with almost twice more high-frequency contacts in 1up12own than 2up1down conformations (see Table 1).

**Table 1:**
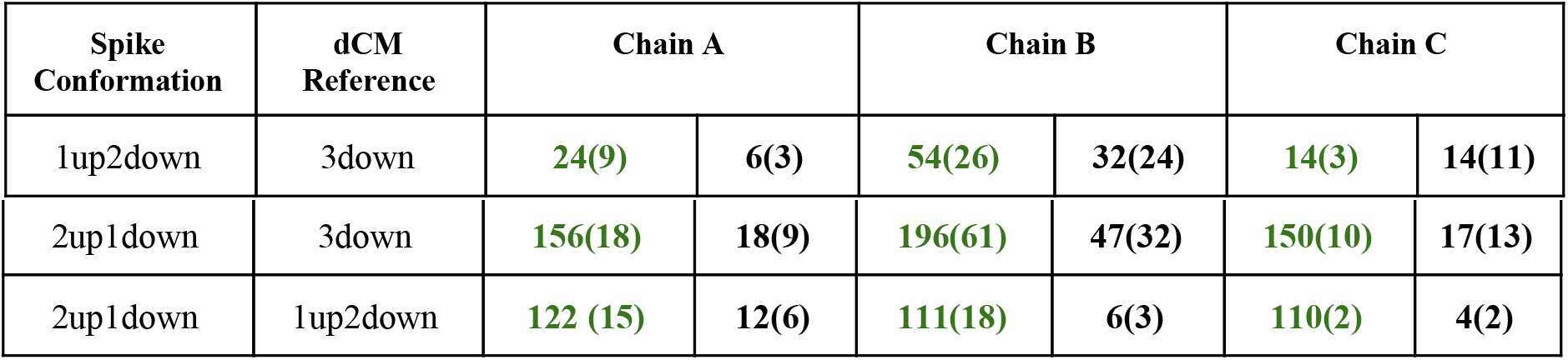
Differential contact map analysis of the spike protein calculated for every single protomer A, B, and C. We show the total number of destabilizing/stabilizing residues in green/black and the corresponding number of intrachain contacts with freq > 0.9 in parentheses.

| Spike Conformation | dCM Reference | Chain A |  | Chain B |  | Chain C |  |
| --- | --- | --- | --- | --- | --- | --- | --- |
| 1up2down | 3down | 24(9) | 6(3) | 54(26) | 32(24) | 14(3) | 14(11) |
| 2up1down | 3down | 156(18) | 18(9) | 196(61) | 47(32) | 150(10) | 17(13) |
| 2up1down | 1up2down | 122 (15) | 12(6) | 111(18) | 6(3) | 110(2) | 4(2) |

**Figure 2:**
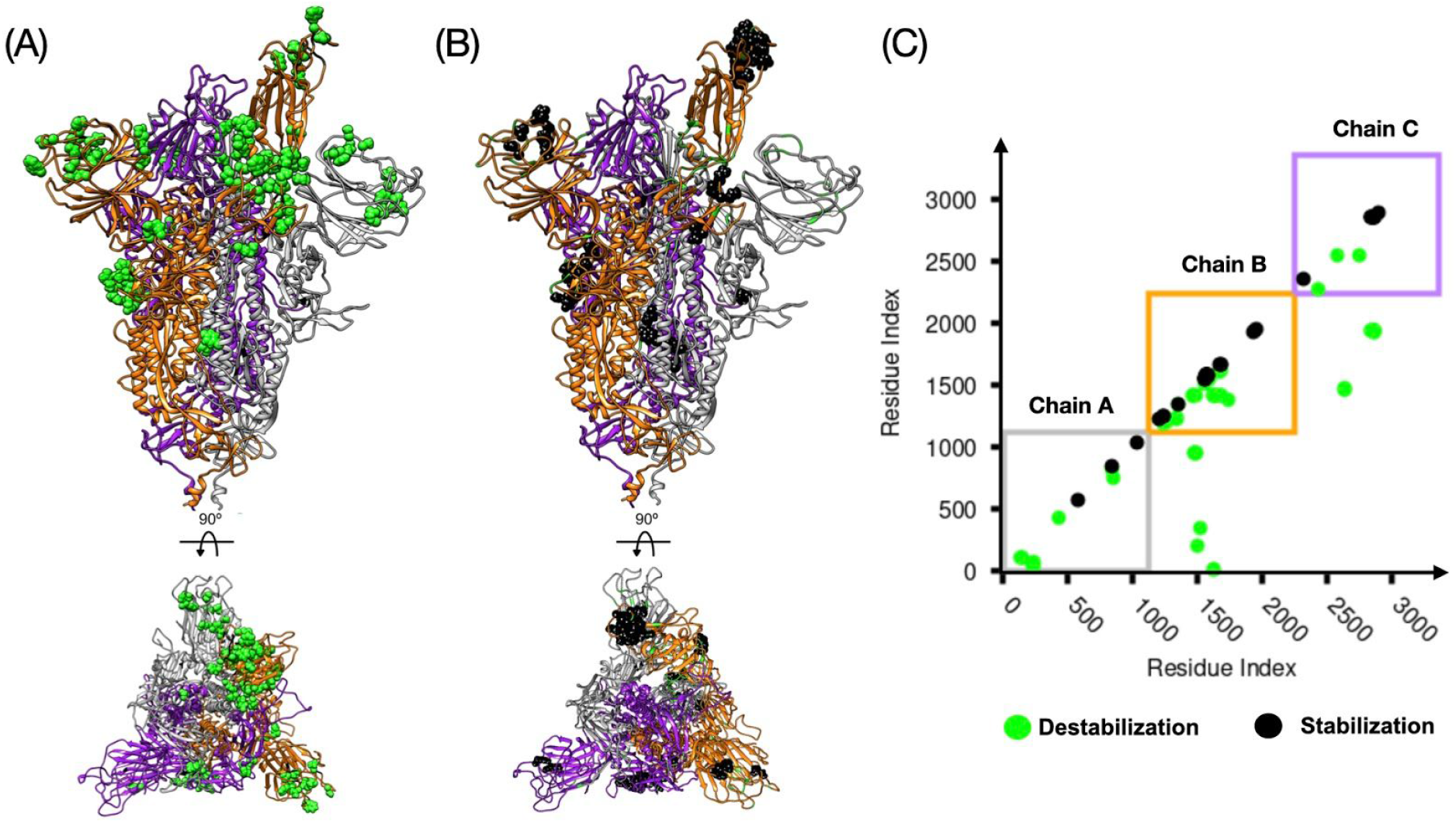
The dCM analysis (f>0.9) of the spike protein with 1up2down RBD. The secondary structure of the chain A, B and C are depicted in gray, yellow and purple respectively. Panel (A) shows the structure of the homotrimer and in green the amino acid residues that destabilize the spike protein in this conformation. The bottom of panel A shows the same structure rotated by 90° as indicated, showing that most of the destabilizing residues are positioned between the RBD (in up position) from chain B and the RBD and NTD from chain A. Panel (B) shows the stabilizing residues in black. Panel (C) shows the plot of the contact map. The squares follow the color convention for each protomer and each dot represents a high-frequency contact (i.e. green=destabilizing and black=stabilizing).

**Figure 3:**
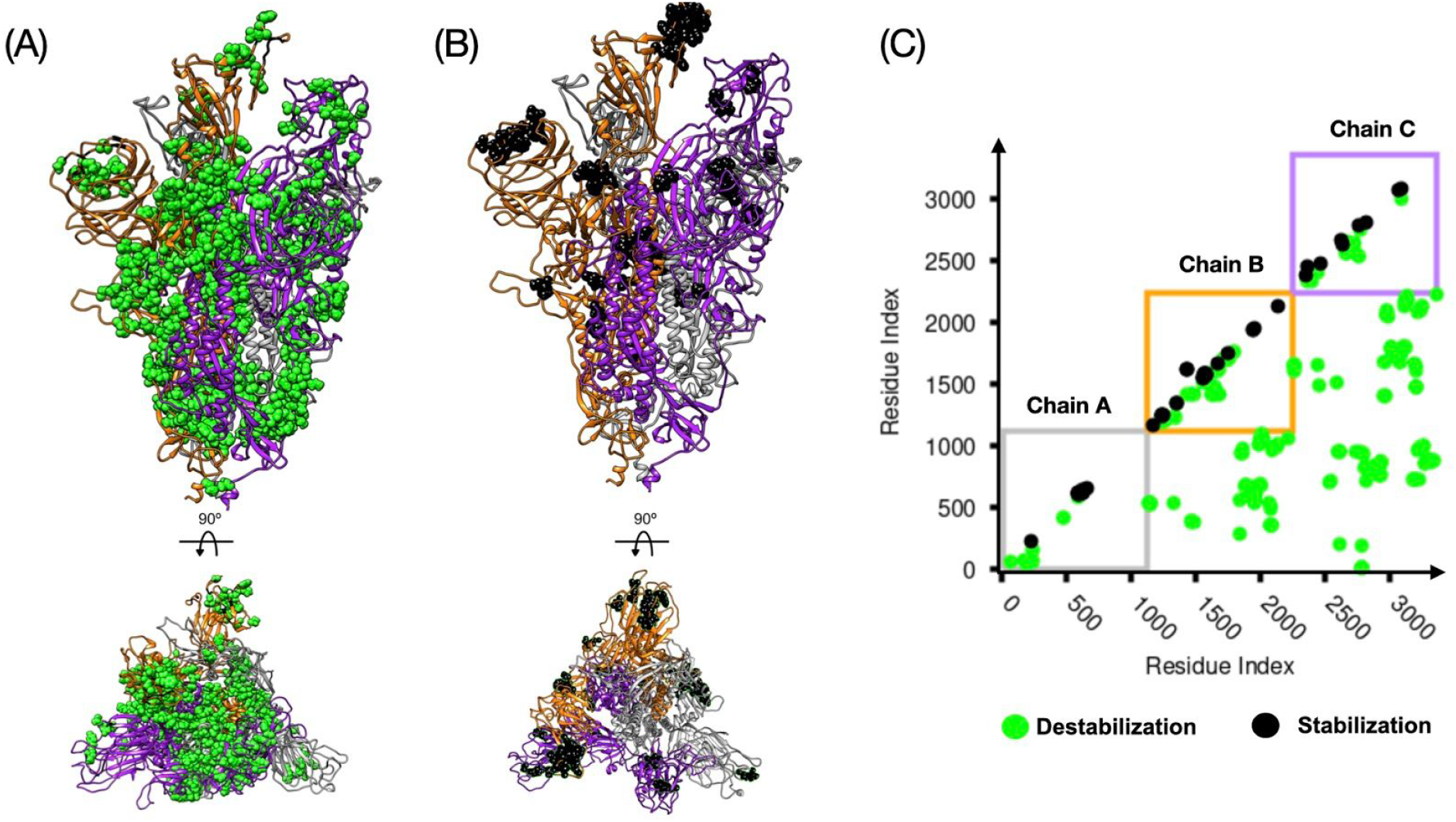
The same analysis as in Figure 2 but for the spike protein with 2up1down RBD conformation.

### Insight into the RBD SARS-CoV-2 conformations

In Fig. 4A we show the RBD in chain B in contact with the RBD in chain C and NTD in chain A in closed conformation. Based on the analysis of the high-frequency contacts (freq >0.9) we find out 25 stabilizing residues in down conformation making 83 contacts that are key for the 1up2down case forming 40 contacts (see Fig. 4B). Such conformational transition emerges through the rearrangement of certain residues located at the bottom of the RBD, close to the hinge region (H519-S530) in chain B that change conformation and breaks 10 high-frequency contacts between hinge region and S2 in chain A in order to transit to the new position. By considering a range of energy values for native contacts in protein, i.e. 1.0-1.5 kcal/mol. The energy associated with the hinge regions in the 3down to 1up2down transition will be in the range of 10-15 kcal/mol. The identification of the residues that create contacts to stabilize a conformation is of great importance for novel therapeutic methods that can target binding to enhance or disrupt the established contacts for a given purpose. For instance, the work by Toelzer, C. *et al*.^21^ has unveiled a free fatty acid (FFA) binding a hydrophobic pocket that locks the structure of the spike in the closed state (see Figure 4C). Such a process is achieved by establishing additional contacts between two nearest RBDs through the FFA. Indeed, such a process is possible due to few pre-existing native contacts in the vicinity of those two RBDs. Those contacts engage the RBD segment (R408-Q409) in chain B and a gating helix (Y365-Y369) in the other RBD in chain A. The latter motif is crucial for the transition from apo conformation to FFA conjugated with protein that has been reported to lock the RBD in the closed state. Our analysis shows the presence of other stabilizing contacts among the gating helix in a given chain and the next chain namely, Q409-Y369, Q414-Y369, T415-Y369, G416-Y369, K417-N370, K417-Y369 and D420-Y369. Due to the symmetry in the closed state, the set of contacts are similarly distributed among all chains. In a recent study by Casalino *et al*.^23^ has been elucidated the stabilization of the open conformation by two additional N-glycan at positions N165 and N234, that are located at the NTD in chain A and according to all-atom MD simulations they modulate the RBD conformational dynamics. Our analysis further shows the stabilization of the RBD in chain B in up conformation through residues located at the NTD in chain A, which are close to N165. Here we get T167(A)-R357(B), F168(A)-R357(B), E169(A)-R357(B), and close to N234 we have P230(A)-T523(B), P230(A)-A520(B) and P230(A)-P521(B). In addition, we also found few residues in RBD in chain A interacting with the RBD in chain C, both considered in down conformation. Those contacts are K417(A)-Y369(C), R403(A)-A372(C), T415(A)-P384(C), K417(A)-N370(C) and L455(A)-N370(C) and more interestingly, we observe an interaction between RBD in up conformation and the closer RBD in down position by establishing contacts such as F377(B)-F486(C), T385(B)-K458(C), S375(B)-F486(C), S383(B)-F456(C), T376(B)-F486(C), T385(B)-F456(C) and F374(B)-F486(C).

**Figure 4:**
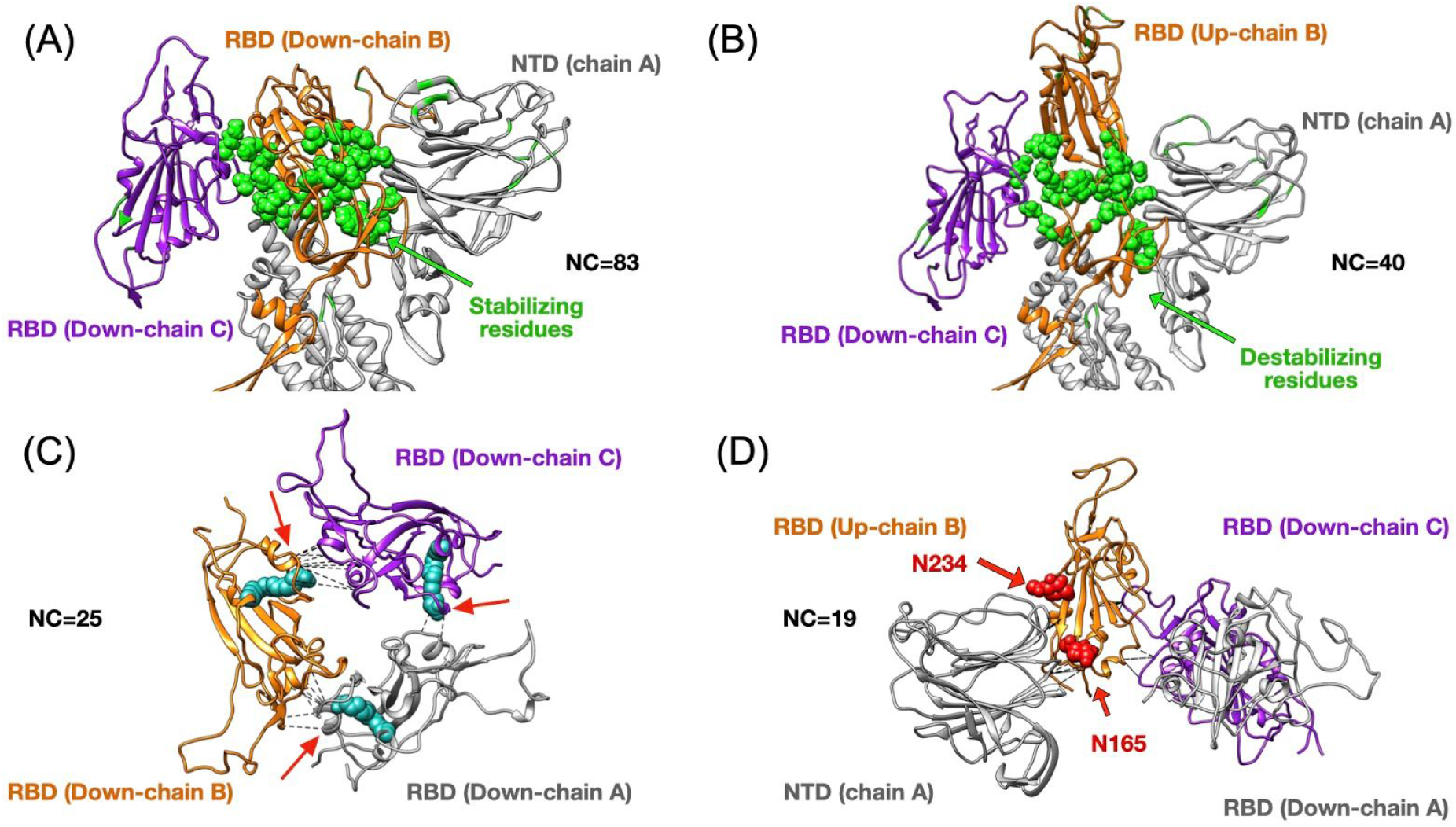
Top panels show changes in terms of stability between the closed (3down) and open (1up2down) states. Panel (A) shows the case of the RBD in chain B stabilized by neighboring protein domains such as the RBD and NTD from chain C and A respectively. The stabilizing 25 residues are highlighted in green and they establish several high-frequency native contacts (NC) equal to 83. Panel (B) shows the same set of residues from panel (A) that are destabilized in the open state forming 40 contacts. Panel (C) shows the stabilization due to 25 contacts between all RBDs in the closed state. The structure of the LA (in cyan) has been superimposed onto our results as it was shown to lock the closed state by forming contacts between two adjacent RBDs. Panel (D) depicts the RBD in up conformation that has been stabilized by 19 contacts formed between the NTD and two other RBDs. The positions of two N-glycans that assist structurally by making extensive interaction with the RBD in the up conformation have been superimposed onto our structure. We highlight the residue contacts that are responsible for stabilization by dashed black lines.

### Energetic calculations of the spike protein conformations

The role of electrostatic interactions in the spike proteins have been calculated by the well known Poisson-Boltzmann (PB) method^51^. It is composed by two contributions, namely, the solvation energies and a complementary term originated by coulomb interactions. The solvation energies which have been calculated by capturing the spike proteins structure from the MD trajectory^40^, and they depend on the solvent-excluded surface (SES) and the partial charges of the solute. A second contribution arises from the point charges distribution in the solute, given by the Coulomb energy, (see the Methods section for the theoretical background). The PB scheme has several advantages, when tackling large biggish protein assemblies of different structures and trajectories provided, as it is the case of the coronavirus spike proteins PDBs. One of the key advantages is the computational feasibility when tackling protein assemblies in the order of 100 thousands of atoms in just an hour^52^.

In Table 2, we have calculated the PB-energy differences between the spike proteins with at least one RBD up and with the closed spike configuration as a reference. Assuming physiological conditions given by a pH of 7 and ionic-strength of 150mM, we observed that the ground energy state is the spike conformation with 3down. In addition, the most favorable open conformation is given by the 1up2down conformation, followed by the 2up1down conformation. Interestingly the energy barrier to reach a 2up1down conformation seems to have an energetic penalty much higher compared to as the first transition, i.e. 3down to 1up2down. Here, we remark that PB-energies provide a static picture of the solvation and Coulomb energies at a certain point of the MD simulations trajectory. Table 2, shows the calculation of the ΔG of energy for 3 MD snapshots corresponding to a tuned mesh refinement (4.16 elements/ Å^2^) set of the three system ending conformations^40^.

**Table 2:** ΔG calculated by the PB method at a pH of 7 and a ionic-strength of 150mM.

| Spike Conformation<br>“C” | Spike Conformation<br>Reference “R” | $\Delta(C-R)$ PB energies<br>[kcal/mol] |
| --- | --- | --- |
| 1up2down | 3down | 10.438 |
| 2up1down | 3down | 32.462 |

The PB energy calculation can be considered as an initial overall energetic comparison between the different spike protein conformations, which also goes in line with the contact map stability analysis.

However given the relevance of the recognition process, we are not providing a final assessment of the ΔGs. In fact, we suggest extending this study and comparing results between different free-energy calculation methods^53^ applied to the spike protein including a reconstruction of the envelope membrane. The inclusion of a reconstructed membrane may provide further insight into the large and flexible conformational space of the spike proteins in a more robust and accurate manner.

## CONCLUSIONS

In the current COVID-19 pandemic the development of novel diagnostics and antiviral therapies is of high priority. We expect our structural and energetic studies can provide additional information regarding the space of interactions mapped by certain key residues that are crucial for stabilization of the spike protein during an apparent dynamic equilibrium mediated by transitions from close to open conformations prior to ACE2 recognition. In particular we found several high-frequency contacts formed between the NTD (in chain A) and RBD (in chain B) that are responsible for the local conformational stability, and our data suggest that these high-frequency contacts play a role during the transition from closed to open state. Our analysis shows that this transition occurs at the energetic cost of breaking very high-frequency contacts between the RBD hinge and the S2 region in chain B and A respectively. The rearrangement of those residues has an energetic cost in the range of 10-15 kcal/mol which is consistent with the PB analysis that quantifies the change in free energy on the order of 10.4 kcal/mol for 3down to 1up2down. Our studies also show the large energetic cost required to transit from closed to 2up1down conformation (~ 30 kcal/mol) in the absence of the cellular ACE2. This results indicates the propensity of the spike protein to be found likely in the 1up2down conformation prior to interacting with the cell surface. Further studies based on single molecule force spectroscopy can help to differentiate spike protein conformation according to their mechanical properties (e.g Young modulus).

In perspective, we aim to provide enough information about possible target sites to destabilize the spike conformations of the closed and open state (i.e. 1up2down or 2up1down). One possibility is that our work can be combined with other studies on the binding of natural compounds derived from plant sources^54^. In this regard, several herbals derivatives having hepatoprotective, anxiolytic, antidepressant, nootropic, antimicrobial, anti-inflammatory, antioxidant, anti-stress, anticonvulsant, cardio-protective, antitumor, anti-genotoxic, anti-Parkinson and immunomodulatory properties can be used to target inhibition of spike protein conformation. Recently, it has been approved to use them as the alternative antiviral inhibitor for COVID-19 patient treatment^28,29^. In view of the above considerations, in the future, we plan to investigate and determine the efficacy of the potential herbal candidates along with FDA approved drugs (e.g. Remdesivir, Lopinavir, favipiravir and Hydroxychloroquine)^4^ against SARS-CoV-2 spike protein by destabilizing the RBD interactions with ACE2. This can be accomplished by combining in-silico docking and molecular dynamics simulation in order to determine the destabilizing residues and how their contacts with the RBD are broken upon ligand binding. From this work, we will be able to propose possible computationally promising compounds that can be further probed experimentally.

## Author contribution

A. B. P, and R. A. M designed the research; all authors performed the calculation, analyzed data, and wrote the paper; and A. B. P supervised the research.

## Acknowledgments

A. B. P. and R. A. M. thank the National Science Centre, Poland, for financial support under grant No. 2017/26/D/NZ1/00466. H. V. G thanks the Slovenian Research Agency for financial support (Funding No. P1-0055). J. L. B. acknowledges the use of the ELSA high-performance computing cluster at The College of New Jersey. This cluster is funded in part by the National Science Foundation under grant numbers OAC-1826915 and OAC-1828163. The authors gratefully acknowledge the computing provided by the Jülich Supercomputing Centre on the supercomputer JURECA at Forschungszentrum Jülich. This research was supported in part by PLGrid Infrastructure. S.B. acknowledges Dirección General de Asuntos del Personal Académico de la Universidad Nacional Autónoma de México (DGAPA-UNAM) for a postdoctoral fellowship.

## Notes

https://zenodo.org/record/4165156#.X53L7Gko8zQ

### Competing Interest Statement

The authors have declared no competing interest.

